# PhenoTrack3D: an automatic high-throughput phenotyping pipeline to track maize organs over time

**DOI:** 10.1101/2022.07.19.500623

**Authors:** Benoit Daviet, Romain Fernandez, Llorenç Cabrera-Bosquet, Christophe Pradal, Christian Fournier

**Affiliations:** LEPSE, Univ Montpellier, INRAE, Institut Agro, Montpellier, France; CIRAD, UMR AGAP Institut, F-34398 Montpellier, France; UMR AGAP Institut, Univ Montpellier, CIRAD, INRAE, Institut Agro, F-34398 Montpellier, France; Inria & LIRMM, Univ Montpellier, CNRS, Montpellier, France

**Keywords:** high-throughput phenotyping, computer vision, maize, tracking, sequence alignment, plant physiology

## Abstract

**Background:** High-throughput phenotyping platforms allow the study of the form and function of a large number of genotypes subjected to different growing conditions (GxE). A number of image acquisition and processing pipelines have been developed to automate this process, for micro-plots in the field and for individual plants in controlled conditions. Capturing shoot development requires extracting from images both the evolution of the 3D plant architecture as a whole, and a temporal tracking of the growth of its organs.

**Results:** We propose PhenoTrack3D, a new pipeline to extract a 3D+t reconstruction of maize at organ level from plant images. It allows the study of plant architecture and individual organ development over time during the entire growth cycle. PhenoTrack3D improves a former method limited to 3D reconstruction at a single time point [Artzet *et al*., 2019] by (i) a novel stem detection method based on deep-learning and (ii) a new and original multiple sequence alignment method to perform the temporal tracking of ligulated leaves. Our method exploits both the consistent geometry of ligulated leaves over time and the unambiguous topology of the stem axis. Growing leaves are tracked afterwards with a distance-based approach. This pipeline is validated on a challenging dataset of 60 maize hybrids imaged daily from emergence to maturity in the PhenoArch platform (ca. 250,000 images). Stem tip was precisely detected over time (RMSE < 2.1cm). 97.7% and 85.3% of ligulated and growing leaves respectively were assigned to the correct rank after tracking, on 30 plants x 43 dates. The pipeline allowed to extract various development and architecture traits at organ level, with good correlation to manual observations overall, on random subsets of 10 to 355 plants.

**Conclusions:** We developed a novel phenotyping method based on sequence alignment and deep-learning. It allows to characterise automatically and at a high-throughput the development of maize architecture at organ level. It has been validated for hundreds of plants during the entire development cycle, showing its applicability to the GxE analyses of large maize datasets.

## Background

Plant architecture relates to the 3D organisation and form of organs along the plant (i.e. number, shape, size and position) as well as their temporal and topological changes during plant development [Barthélémy and Caraglio, 2007; Bucksch et al., 2017]. Plant architecture and development are of major agronomic importance in cereals such as maize (*Zea mays* L.), determining resource capture and use, and thus plant performance [Stewart et al., 2003; Long et al., 2006; Zhu et al., 2013]. Architectural and developmental traits such as leaf number, leaf angle and leaf size are influenced by both genetic variations and environmental conditions such as light and water availability [Cabrera-Bosquet et al., 2016; Lacube et al., 2017; Perez et al., 2019], and display a large genotype x environment (GxE) interaction. Therefore, understanding the genetic control and the response to environmental cues of these traits at individual plant level is a major challenge in the context of climate change [Murchie et al., 2009; Zhu et al., 2010; Reynolds et al., 2012], and the engineering of agro-ecological practices based on complex plant mixtures.

High-throughput phenotyping (HTP) platforms enable the collection of plant images of a large number of genotypes growing under different environmental conditions on a regular basis [Tardieu et al., 2017; Roitsch et al., 2019]. Such image datasets can then be processed automatically [Minervini et al., 2015] to capture the 3D morphology of plant shoots and extract phenotypic traits [Gibbs et al., 2016]. Among the existing methods, the Phenomenal pipeline [Artzet et al., 2019] proved relevant to process large datasets of multiple species of agronomic interest, providing an end-to-end solution to extract a 3D plant reconstruction from a set of 2D images taken at regular viewpoints around the plant. In the case of maize, this pipeline allows the extraction of architectural traits such as stem height and leaf morphology (e.g. length, insertion height, azimuth). However, it is limited to reconstruct plants at each time point separately. When screening for genetic variability in maize, the identification of leaves by a time-consistent number (the rank of emergence, or leaf rank) is essential to compare the shoot organs occupying similar developmental position (e.g. juvenile versus adult leaves [Poethig, 2013]) between different plants, or to quantify the development using leaf stage. Moreover, the assignment of the same rank to successive segmentations of the same leaf over time is necessary to measure individual leaf growth and their responses to environmental conditions. Since the lowest maize leaves disappear over time due to senescence, it may be difficult to deduce the rank of leaves when considering the plant at a single date [Ledent and Mouraux, 1990].

To overcome this limitation, time-series analysis could be used to group several occurrences of a same leaf in a temporal series of images, in order to get a 3D+t representation of the plant (Fig. 1). While this task relates to multiple object tracking [Luo et al., 2020], this framework cannot be directly applied to maize leaf tracking, since plants undergo major topological and morphological changes over time, with new leaves appearing and growing due to organogenesis, and others collapsing then disappearing during senescence [Li et al., 2013]. Instead, specific leaf tracking methods have been proposed for 2D images of rosette plants from a top view [Aksoy et al., 2015; Dellen et al., 2015; Viaud et al., 2017; Yin et al., 2017]. While a few studies have developed leaf tracking methods on 3D reconstructions (e.g. cotton [Paproki et al., 2012], cucumber [Harmening and Paffenholz, 2021], tomato [Chebrolu et al., 2021] and maize [Bashyam et al., 2021; Chebrolu et al., 2021]), most of them have only been validated on limited datasets that may not reflect actual HTP conditions (thousands of plants, large genetic diversity and growing scenarios). Moreover, these datasets were often limited to young plants which are less challenging to analyse than in later growth stages (emergence of the reproductive organs, more occlusion due to leaf crossings, more frequent disappearance of leaves due to senescence). While these methods offer various solutions for associating leaves with similar geometry (e.g. length, azimuth), they rarely use topology (i.e. the spatial organisation of leaves on the plant [Balduzzi et al., 2017]). Plant topology offers valuable information for leaf tracking since (i) it defines the identity of leaves, as leaves appear from bottom to top along the stem axis on plants such as maize, and (ii) it is redundant over time, thus helping to maintain leaves identity. Plant topology was used in [Bashyam et al., 2021], but without considering leaf morphology. We therefore seek a new leaf tracking method exploiting both leaves morphology and topology, with the objective of handling complex HTP datasets, covering all stages of maize development.

**Fig. 1.**
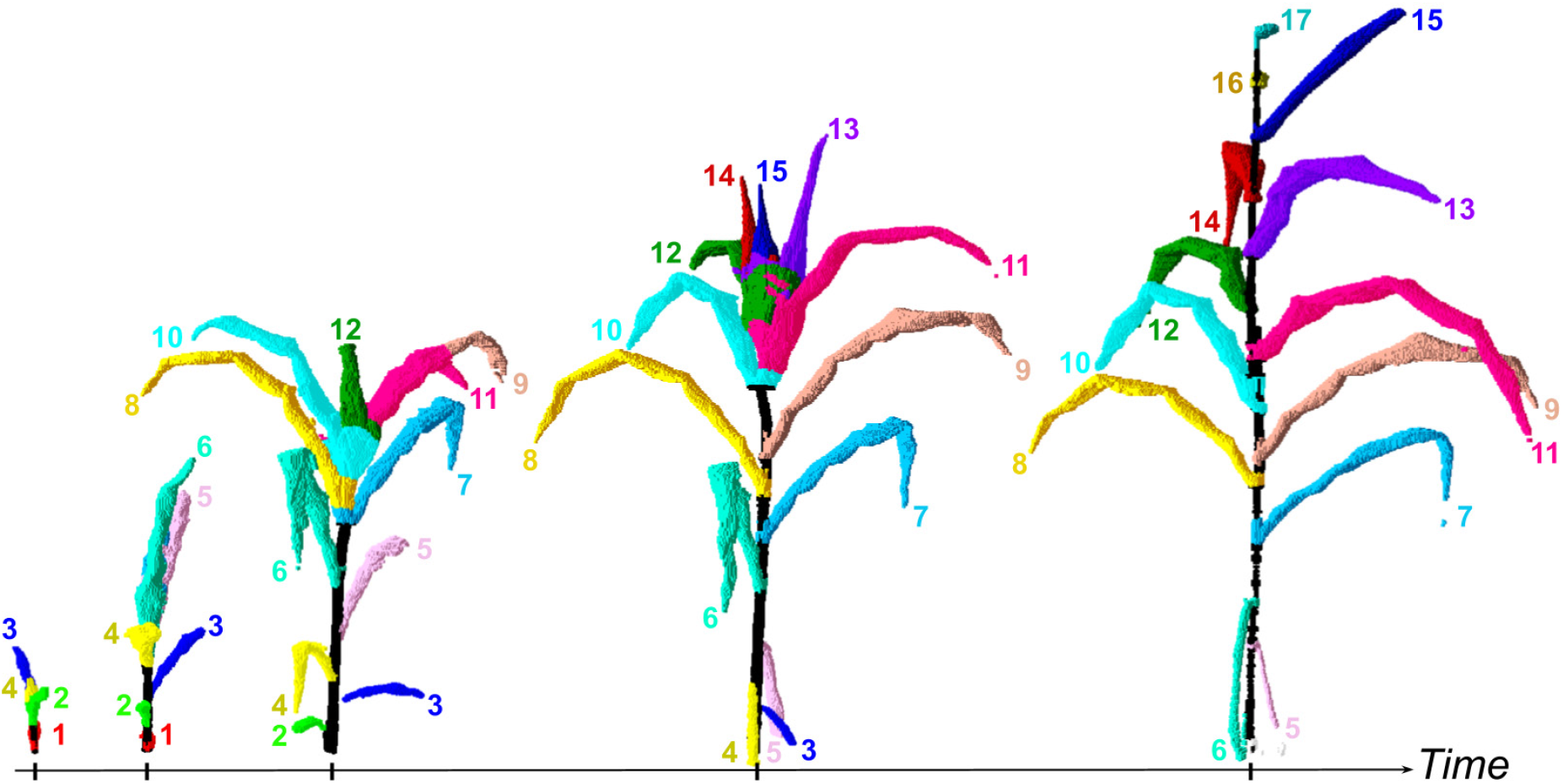
3D+t reconstruction of a maize plant during the whole vegetative development, using the pipeline. The pipeline was run on a time-series of 45 sets of 12 images each, and the outputs are displayed for 5 dates. At each date, the reconstruction is made of a set of voxels segmented into organs. Each segmented leaf is associated to a leaf rank corresponding to its order of emergence, shown by a number and a colour. The stem is displayed in black.

Using first the topological order of leaf ranks along the maize stem, a reconstructed maize plant can be represented at any date by a spatial sequence of leaves, ordered from the bottom to the top of the stem. This sequence is obtained through a segmentation step and may therefore contain artefacts resulting from segmentation errors. For instance, the maize ear can be misidentified as a leaf, or some leaves can be unidentified due to occlusions, leading to extra or missing leaves in the sequence. Some leaves may also emerge or fall from the plant between two successive sequences. From this point of view, comparing two successive leaf sequences is analogous to comparing two homologous genetic sequences, which are similar except for a few extra or missing elements. Sequence alignment algorithms are commonly used to insert gaps in such genetic sequences in order to match their common elements [Batzoglou, 2005]. This framework was redesigned here to match leaves with a similar morphology in successive topological sequences of maize leaves.

In this paper, we propose a novel robust method to address the current issues of maize HTP. Organs are segmented on 3D reconstructed volumes at all time steps. Leaves of the basal mature parts are aligned using a sequence alignment algorithm, then tracked backwards to the upper developing parts of the plant. We apply this method on a challenging image dataset acquired at the PhenoArch platform consisting of a diverse panel of maize genotypes developing from plant emergence to late flowering stage, under various levels of water stress. The quality of the tracking is globally assessed by comparing leaf rank predictions and key dynamical traits to manual annotations. Such organ-level phenotypic traits allow to fully describe the global plant development and architecture, as well as individual leaf growth dynamics.

## Materials and Methods

### Plant material and dataset composition

The pipeline was tested on a dataset from an experiment conducted in 2017 involving a set of 60 commercial maize hybrids representative of breeding history in Europe during the last 60 years. This material covers a wide range of plant architecture, growth and development, leading to an appreciable variability of performances in the field [Welcker et al., 2022]. The experiment was conducted in the PhenoArch phenotyping platform (Fig. 2A) hosted at the M3P (Montpellier Plant Phenotyping Platforms) [Cabrera-Bosquet et al., 2016].

**Fig. 2.**
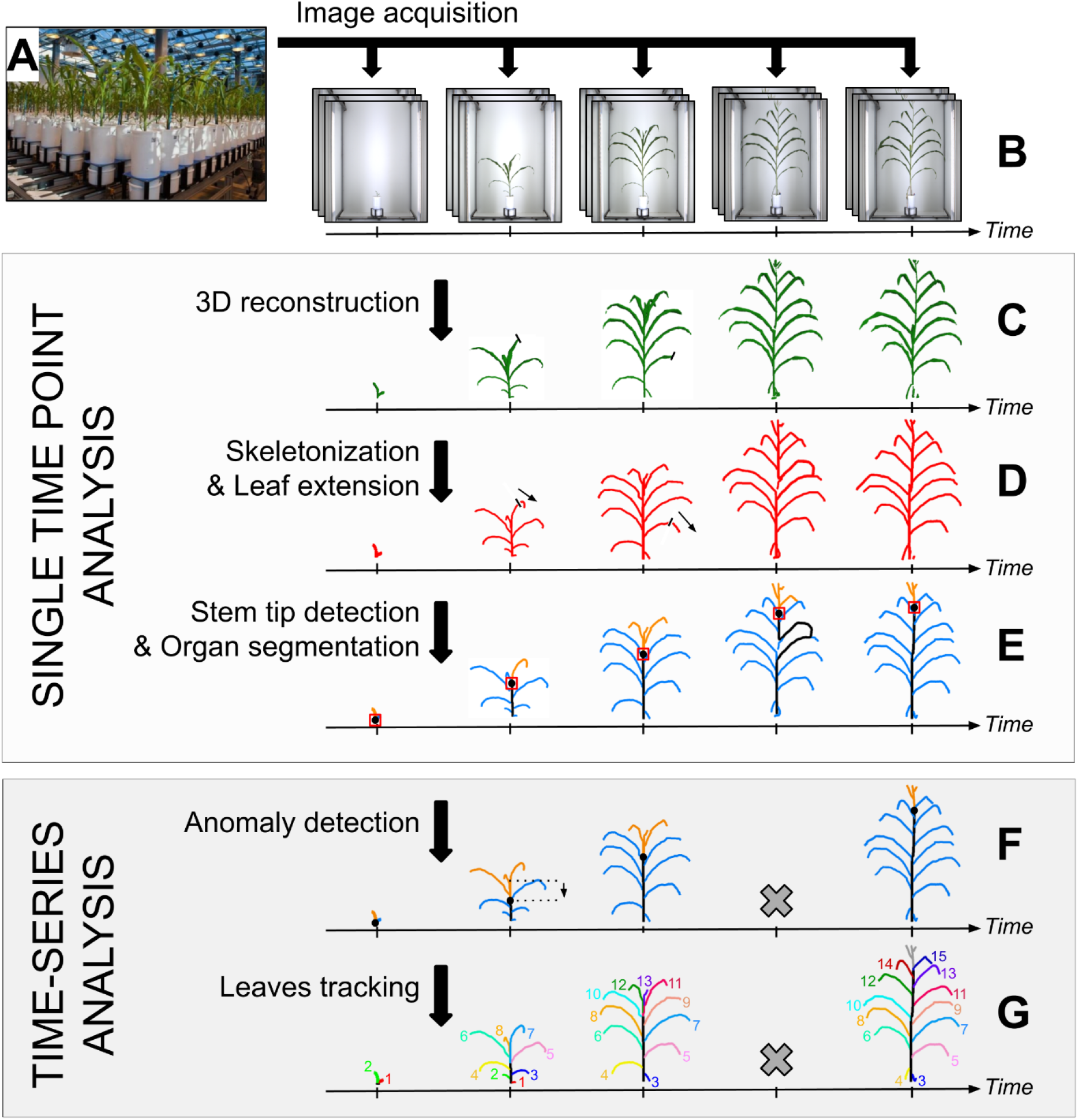
Overview of the 3D+t maize reconstruction pipeline. **A)** Maize plants grown in the PhenoArch high-throughput phenotyping platform. **B)** Daily acquisition of 12 side-view RGB images. **C)** Reconstruction of 3D volumes. **D)** 3D skeletonization of the reconstructions, and extension of the leaf tips. **E)** Detection of the stem tip (red box) using a deep-learning model. This position is used to segment the skeleton into stem (black), ligulated leaves (blue) and growing leaves (orange) organs. **F)** Smoothing of stem height over time and removal of time points with an abnormal stem shape (cross). **G)** Leaf temporal tracking and rank assignment. Each leaf rank corresponds to one colour and one number. **B, C, D, E** are performed independently for each date, while **F, G** use the entire time-series at once.

Briefly, plants were sown in 9L pots filled with a 30:70 (v/v) mixture of a clay and organic compost. Two levels of soil water content were imposed: (i) retention capacity (WW, soil water potential of - 0.05 MPa) and (ii) water deficit (WD, soil water potential of - 0.3 MPa). Each combination of genotype and water treatment was replicated 7 times, 4 with early harvesting (until 12 visible leaves stage, ~40 days after plant emergence) and 3 with late harvesting (until ~55 days after plant emergence, i.e ~15 days after panicle emergence), resulting in a total of 840 plants. Greenhouse temperature was maintained at 25 ± 3 °C during the day and 20 °C during the night. Details of experimental growing conditions can be found at [Brichet et al., 2017].

RGB images (2048 × 2448 pixels) were taken daily for each plant with twelve side views from 30° rotational difference (Fig. 2B), using the imaging units of the PhenoArch platform. Each unit is composed of a cabin involving an RGB camera (Grasshopper3, Point Grey Research, Richmond, BC, Canada) equipped with 12.5–75 mm TV zoom lens (Pentax, Ricoh Imaging, France) and LED illumination (5050–6500 K colour temperature). Images were captured while the plant was rotating at constant rate (20 rpm) using a brushless motor (Rexroth, Germany).

### 3D organ segmentation

The Phenomenal pipeline [Artzet et al., 2019] was applied on our single time-point data to reconstruct 3D volumes of the plants (Fig. 2C). These volumes were skeletonized (Fig. 2D) and individual plant organs (stem, leaves) were segmented (Fig. 2E).

We post-processed the skeleton output by projecting it on the corresponding 2D binary images, to extend the leaf tips which are often shortened during the reconstruction. To that end, binary images extracted with Phenomenal were also skeletonized, and the segments of both skeletons were matched to find an extension path for each 3D segment (Fig. 3).

**Fig. 3.**
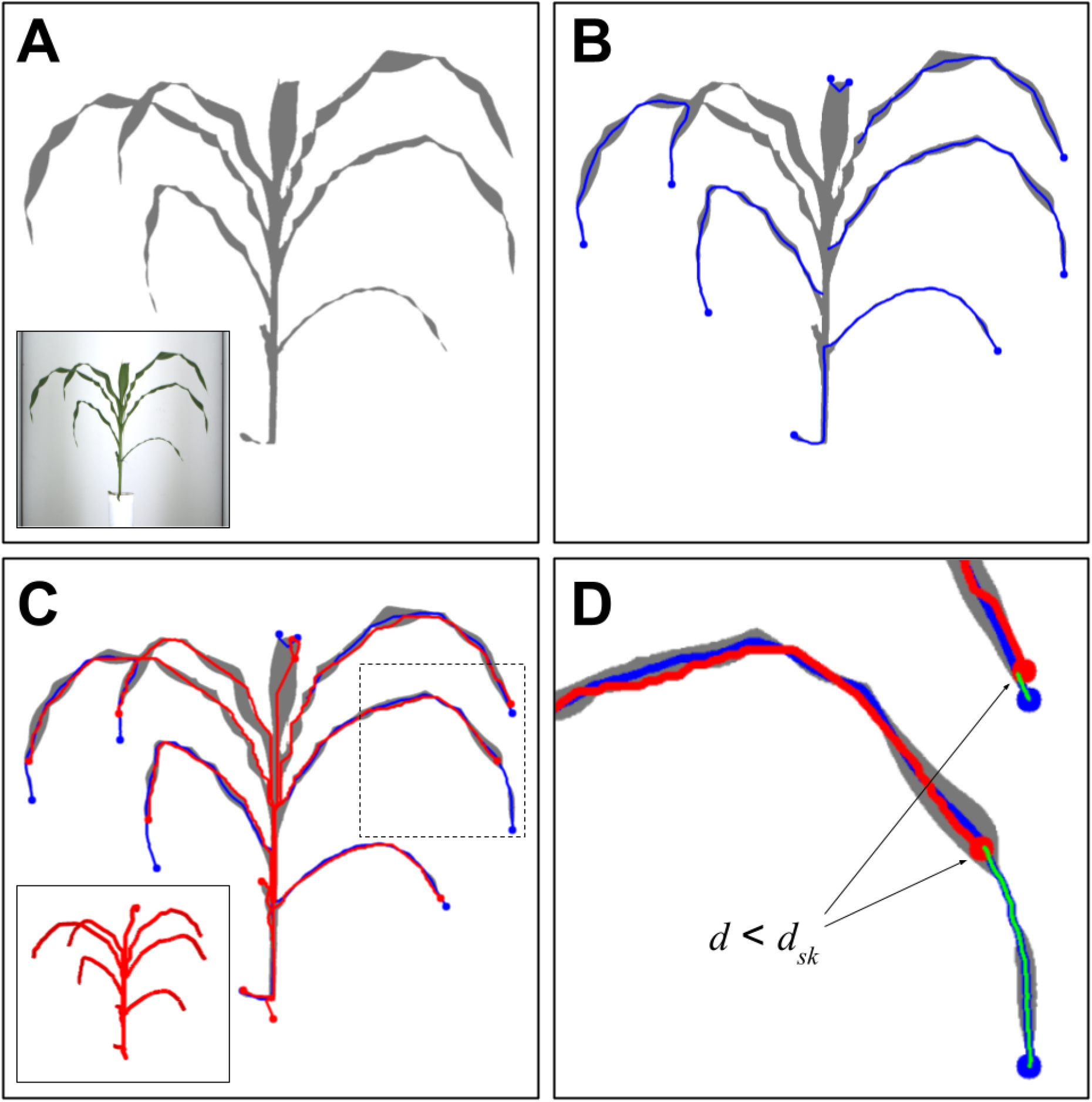
Extension of leaf tips on the 3D skeleton. **A)** Binarization of the RGB image (inset) with Phenomenal [Artzet et al., 2019], **B)** Skeletonization of the binary image, and extraction of skeleton branches having an endpoint (blue lines), **C)** 3D plant skeleton (inset) and reprojection of its branches in the 2D space (red lines). **D)** 3D and 2D branches matching based on a distance threshold *d_sk_* = 30px, and determination of extension paths (green lines).

Finally, we trained an object detection model to detect collars on the 2D images, like in [Zhou et al., 2021]. To that end, we used a YOLOv4 deep-learning model [Bochkovskiy et al., 2020] (see Additional file 1 for training details). The detected collars were used to define the stem height as the highest collar height among the 12 side-view images. The leaves with their insertion point below on the stem were defined as ligulated leaves (Fig. 4).

**Fig. 4.**
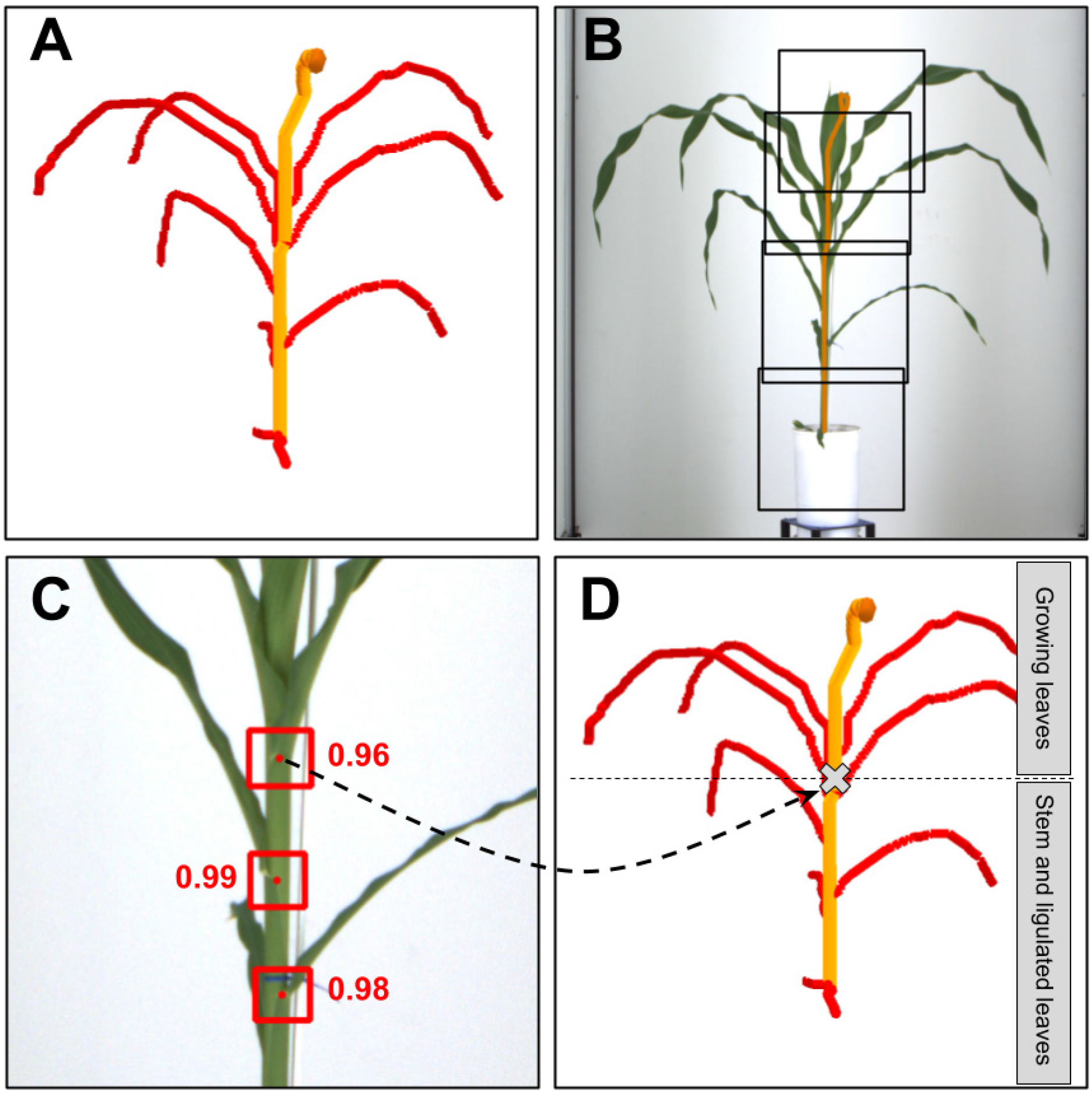
Deep-learning-based collar detection for organ segmentation. **A)** Identification of the stem path (orange line) on the 3D skeleton, **B)** reprojection of the stem path (orange line) on one of the corresponding RGB images and extraction of 416×416 sub-images along the stem path, **C)** Collar detection on a sub-image (square: predicted bounding box, point: centre of the box, value: prediction score) **D)** Projection of the highest detected collars point among all sub-images from all 12 side viewpoints on the 3D skeleton (grey cross). This point is then used to segment skeleton branches in stem, ligulated leaves, and growing leaves.

From this point, each plant is represented by a time-series of 3D observations, from which stem height is smoothed over time, and time points with abnormal stem shapes are removed (Fig. 2F).

### Time-lapse tracking of 3D organs

We track the ligulated leaves using a multiple sequence alignment algorithm. First, at each time point *t* we build a leaf sequence *P_t_* composed of feature vectors representing key characteristics of the segmented ligulated leaves. From this representation, we define a cost function for comparing leaves pairwise, and derive a sequence alignment algorithm to establish the correspondences between two successive leaves sequences. We then extend this procedure to determine leaf tracks along the whole time-course. Each organ track is then identified by a leaf rank, which corresponds both to its order of emergence and its location along the stem. Finally, we track backwards these leaves from the moment of their ligulation to the moment of their emergence at the top of the stem and get the whole 3D+t reconstruction.

#### a) Pairwise sequence alignment

The detected ligulated leaves of a date *t* are ordered from the bottom to the top of the plant, i.e. by ascending leaf rank, in a leaf sequence *S_t_*. The alignment of two sequences can be defined as a set of gap placements at the beginning, end, or between elements of these sequences, resulting in two new sequences of the same length with no gaps facing each other [Batzoglou, 2005]. Pairwise sequence alignment algorithms are designed to identify the alignment that optimises an alignment score between two sequences, often defined as the sum of the scores associated with each pair of matched elements, plus gap penalties [Edgar and Batzoglou, 2006].

Each ligulated leaf is described using a feature vector computed from geometrical features that are assumed to be constant over time: insertion height *h (*mm), length *l (*mm*) a*nd azimuth *α* (*α* ∈ [-1, 1]). These features are extracted from the segmented leaves [Artzet et al., 2019] and concatenated into a feature vector 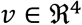 summarizing the leaf morphology:

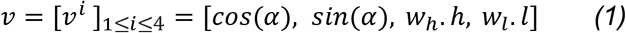

Weights *w_h_* = 0.03 and *w_l_* = 0.004 were fine-tuned to scale features and adjust their relative importance. Each pair of matched leaves is associated to a cost *c_vv_* equal to the euclidean distance between the feature vectors *v_1_* and *v_2_* of those two leaves:

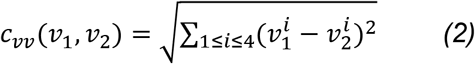

A gap penalty parameter *g* is used to penalise the addition of each of the *n* gaps placed in the alignment of sequences *S*_*t*1_ and *S*_*t*2_ We define *g* from 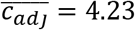, the average value of *c_vv_(v_i_, v_j_*) for all couples of topologically adjacent leaves *i* and *j* in the sequences *S_t_* of our dataset, and *w_gap_* = 3 a weight:

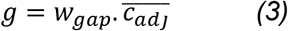

Using a different parameter value for the terminal gaps has proven effective in aligning sequences of different lengths [Edgar, 2004]. A weight *w_tml_* = 0.2 is therefore used to lower the penalty of each of the *n_tml_* terminal gaps, since leaves are expected to appear and disappear successively over time. The optimal alignment between *S*_*t*1_ and *S*_*t*2_ is then defined as the one that minimises the global alignment cost *C*. We define *I* as the set of indexes of sequences *S*_*t*1_ and *S*_*t*2_ with a match (i.e. without gap). With *v_i_* and *v’*_*i*_ the vectors associated respectively with the *i*-th elements of the sequences *P*_*t*1_ and *P*_*t*2_

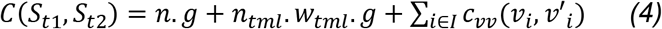

To find the optimal alignment between *S*_*t*1_ and *S*_*t*2_, the Needleman-Wunsch (NW) algorithm [Needleman and Wunsch, 1970] has been adapted to consider the terminal gap weight. This algorithm is based on dynamic programming and guarantees an optimal solution for pairwise alignment.

#### b) Multiple sequence alignment

The initial ordering of each sequence gives a relative leaf rank to each leaf that may differ from their absolute rank, due to the fall of senescent leaves during plant development or due to segmentation errors (Fig. 5A). The alignment of all the leaves sequences in the time-series is achieved using a progressive method [Batzoglou, 2005] to perform the multiple alignment task. It consists in a succession of pairwise alignments of the sequences *S_t_* using the NW algorithm, in ascending temporal order. A profile is defined as an alignment of several sequences treated as a unique sequence of columns [Thompson et al., 1994]. Let *Ω*_1 –>*k*_ be the profile constituted of the alignment of the first *k* sequences in the time-series (*S*)_*k*=1…*T*_, each sequence containing *n* elements after the addition of possible gaps. At each time point *t* >1, the sequence *S_t_* is aligned with the profile *Ω*_1 –> *t*–1_, resulting in a new profile *Ω*_1 –> *t*_. Aligning *S_t_* with *Ω*_1 –> *t*–1_ requires a sequence-profile cost function *c* that we adapt from *c_vv_* [Edgar and Sjölander, 2004]. Let ω be a column of *Ω*_1 –> *t*–1_ of length *t* – 1 containing *t* – 1 – *k* gaps, and *k* leaves associated to the feature vectors {*v*_1_,…, *v_k_*}. Let *v*’ be the feature vector associated with a leaf observation present in the sequence to align *S_t_*. Then the cost *c* of the match between ω and *v*’ is defined as:

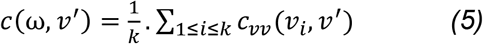

**Fig. 5.**
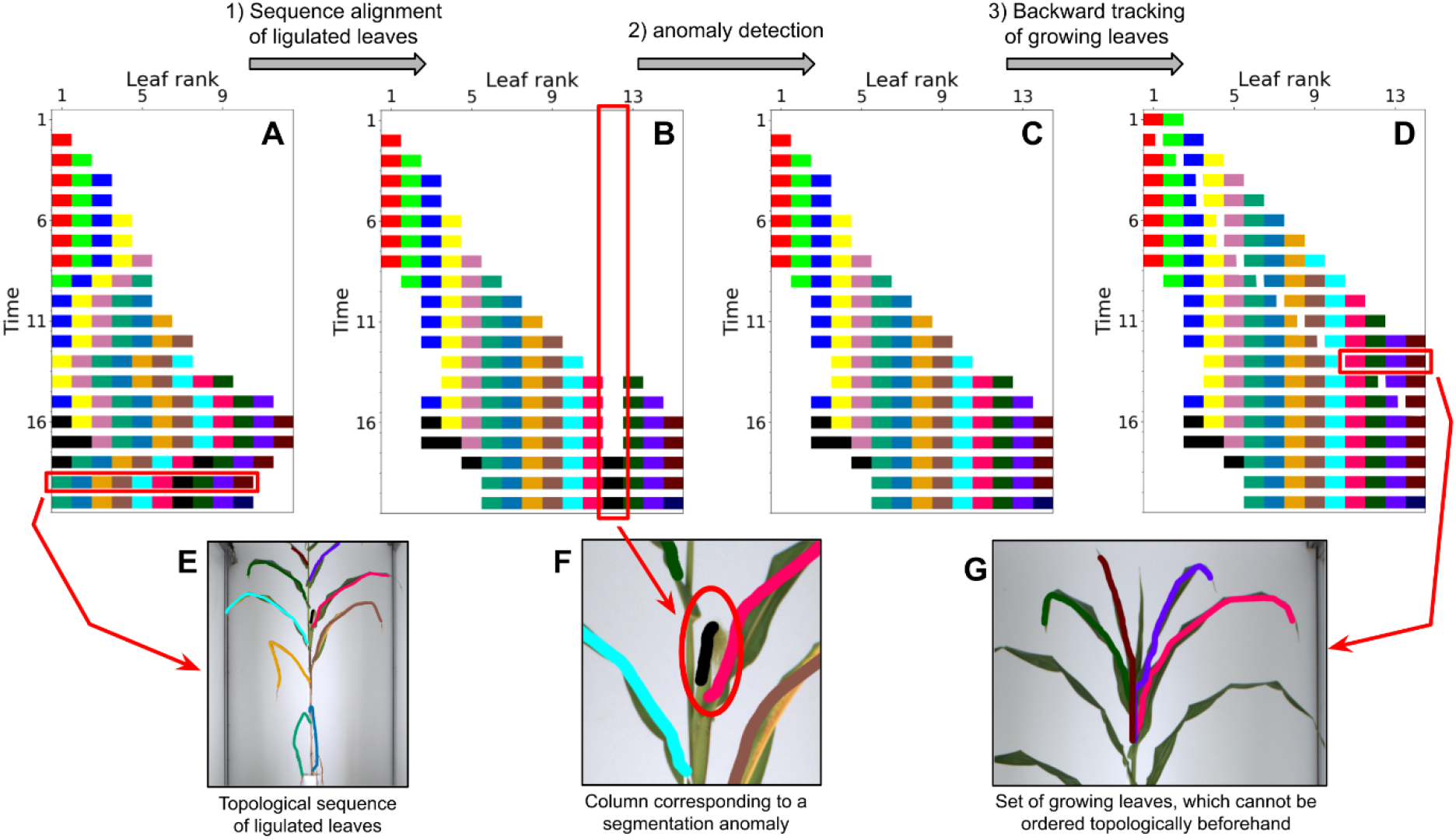
Leaf rank assignment in the time-series of 3D segmented maize plants during leaves tracking. Each segmented leaf is represented by a rectangle, coloured according to its ground-truth rank (black = segmentation anomaly), and positioned according to its observation time and predicted rank. **A)** initialization of rank assignment by ordering the ligulated leaves topologically, **B)** rank assignment after sequence alignment, **C)** rank assignment after removing abnormal columns, **D)** final rank assignment after adding growing leaves. **E, F, G)** projection of some segmented leaf polylines on their corresponding 2D source image. Only one third of the sequences in the time-series are represented for visibility.

Sequences are aligned progressively until obtaining the final profile *Ω*_1 –> *T*_ that aligns all the sequences in (*S*)_*t*=..*T*_, yielding a first estimation of the leaf ranks (Fig. 5B).

Finally, each tracked leaf, except the first and last, is deleted if *n_k_* < (*n*_*k*−1_ + *n*_*k*+1_)/4, *n_k_* being the number of times that the *k^th^* leaf appears in the time-series, to cope with possible segmentation errors (Fig. 5C)

#### c) Backwards tracking of growing leaves from ligulated ones

The final tracking step consists of predicting the rank of the detected growing leaves (Fig. 5D). Unlike ligulated leaves, growing leaves cannot be ordered a priori by their topology since they all emerge from the same point, and their geometry is not constant. However, it can be assumed that the shape, orientation and position of a leaf evolves smoothly over time during its growth phase and leaves can be tracked backwards by associating leaf observations sharing a similar shape, from the ligulated stage to the leaf emergence. To that end, a metric *D* is used to quantify the dissimilarity of two growing leaves, based on the distance between their central line, given as a 3D polyline. Each 3D polyline is converted to a set of *n* = 20 points [*pl*_1_,…, *pl_n_*] regularly spaced along the polyline. With *d*(*p*_1_, *p*_2_) the euclidean distance between two points, we define the distance between two polylines such as:

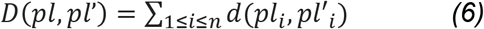

The algorithm tracks backwards the growth trajectory of each leaf, starting from rank 1, and finishing by the last rank. For each processed leaf, the algorithm works iteratively, from the starting point (i.e. when the leaf is ligulated) to the ending point (i.e. at leaf emergence). At each tracking step the algorithm computes the metric *D* between the last associated polyline and every remaining non-ligulated leaves, and selects the minimum.

### Computation of phenotypic traits

Various phenotypic traits are extracted from the 3D+t plant reconstruction, to quantify (i) leaf profiles, (ii) plant development, (iii) individual leaf development.

**Leaf profile** *x_pf_* describes the architecture of the leaves as a function of their rank, regarding a morphological variable *x* (e.g. leaf length, insertion height, azimuth, etc.). For each rank r, *x_pf_*(r) represents the value of *x* for the leaf r once it has reached ligulation. *x_pf_*(r) is calculated as the median of the values of *x* associated with the ligulated leaves along the time course. Leaf observations exceeding 20 day_20°C_ after ligulation are not considered. Here we consider the case of leaf length profile *l_pf_* and leaf insertion height profile *h_pf_*.

**Plant development** is quantified at any date through the following traits:

-Stem height *h_s_* corresponds to the height of the highest collar and is directly extracted from the plant reconstruction.
-Visible leaf stage *n_vis_* corresponds to the rank of the latest emerging leaf. Let *r_vis_*(t) be the maximum rank among observed leaves at date *t*, and *t_med_*(r) the median of time points *t* where *r_vis_*(t) = *r*. We define the emergence timing *t_vis_* of the *r*-th leaf such as:

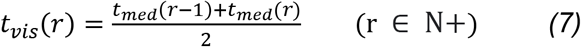

We deduce *n_vis_* such that *n_vis_*(*t*) = *t_vis_*^−1^(*t*) for each emergence timing *t*. Finally, *n_vis_* is extended to any value of *t* by linear interpolation.
-Ligulated leaf stage *n_lig_* corresponds to the rank of the last ligulated leaf. Using the same method as for *n_vis_*, it is deduced from ligulation timing *t_lig_* such as:

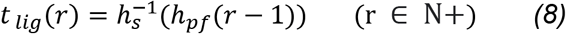

Here, a piecewise constant interpolation is used to restrict *n_lig_* to integer values.

**Leaf growth** is given by the successive length value of the observed *r*-th leaf until ligulation.

### Validation with manual measurements

Ground-truth data was manually collected on a randomly selected subset of plants with late harvesting. This validation data was then used to evaluate tracking performance, and the accuracy of the phenotypic traits obtained with the pipeline.

Leaf ranks were annotated on 30 plants at each time point using the images (10980 annotations). Segmented leaves corresponding to artefacts (e.g. the ear of maize) were not annotated.

A subset of 10 plants were randomly selected for manual measurements on images. Leaf lengths and leaf insertion heights were measured at the ligulation stage for all ranks (113 and 173 annotations respectively). Additional leaf lengths measurements were also performed for leaf ranks 6 and 9 during their whole growth phase (234 annotations). Stem height was annotated for all time points (369 annotations).

Ligulated and visible leaf stages were measured in the greenhouse on the 355 plants with late harvesting, with an average of 7 time points per plant (2289 and 1891 annotations respectively). Ligulated leaf stage is given by an integer, while visible leaf stage is given by a real number: for example, a value of 7.4 means that the last visible leaf is of rank 7, and has reached 40% of its growth.

The phenotypic traits were compared with ground-truth observation using the following metrics: bias, root-mean-square error (RMSE), mean absolute percentage error (MAPE) and coefficient of determination (R^2^).

The pipeline conception and the data analysis were performed with Python.

## Results

### A robust and consistent alignment of segmented plants over time

The full pipeline was run on the whole late harvesting dataset, (355 plants). Each plant was observed on an average of 43 time points, from plant emergence until 3 days after panicle deployment, making a total of 237,600 images analysed. Leaf tracking output is illustrated in video in Additional file 2. 1.7% of time points were automatically discarded at the anomaly detection step (Fig. 2F), due to abnormally shaped stems (Additional file 3). The global quality of the pipeline was assessed by leaf rank assignment accuracy, which is defined as the percentage of exact matches between the predicted rank of segmented leaves and manually annotated ground-truth ranks (observation). This metric was evaluated separately depending on whether leaves were identified as ligulated or growing, since their tracking relies on different algorithms (Table 1).

**Table 1.** Evaluation of the performance of the maize leaf tracking algorithm. Leaf rank assignment accuracy is evaluated on 30 different plants on a total of 10980 leaves. Leaf rank assignment accuracy is the percentage of non-artefact segmented leaves whose predicted rank matches ground-truth rank. MAE is the mean absolute error, and this metric was only computed among wrong predictions. n is the number of leaves considered. These metrics are presented separately for ligulated and growing leaves.

| Type of segmented leaf | Leaf rank assignment accuracy (%) | MAE among wrong predictions | n |
| --- | --- | --- | --- |
| Ligulated leaf | 97.7 | 1.07 | 6540 |
| Growing leaf | 85.3 | 1.80 | 4440 |

For ligulated leaves, rank assignment accuracy showed a median value of 98.8% per plant, with a minimum of 90.8%, resulting in a high overall accuracy (97.7%, Table 1). Most of the errors occurred for the lower and upper ranks (Fig. 6, blue line), but the resulting rank error did not exceed 1 most of the time (MAE = 1.07 among wrong predictions, Table 1). Errors in ranks 1-3 could be partly removed and errors in ranks 4-5 almost completely removed by putting aside the leaves that grow older than 20 day_20°C_ (Fig. 6; blue dotted line). Such old leaves are harder to track, as their morphology tends to change excessively when senescing. The remaining errors in ranks 1-3 were probably due to segmentation issues. For instance, the leaf 1 is sometimes not segmented in the 3D reconstruction because of its small size (data not shown). About half of the errors in the upper ranks (10 and more) could be avoided by manually removing the dates where the maize ear was misidentified as a leaf during the segmentation process (Fig. 6; blue dotted line), demonstrating the importance of a correct identification of this organ. The remaining errors in the upper ranks might be due to the increasing complexity of maize architecture during its development (longer leaves, more occlusions due to leaf crossings), and because these late emerging leaves are observed fewer times, making their identification more difficult.

**Fig. 6.**
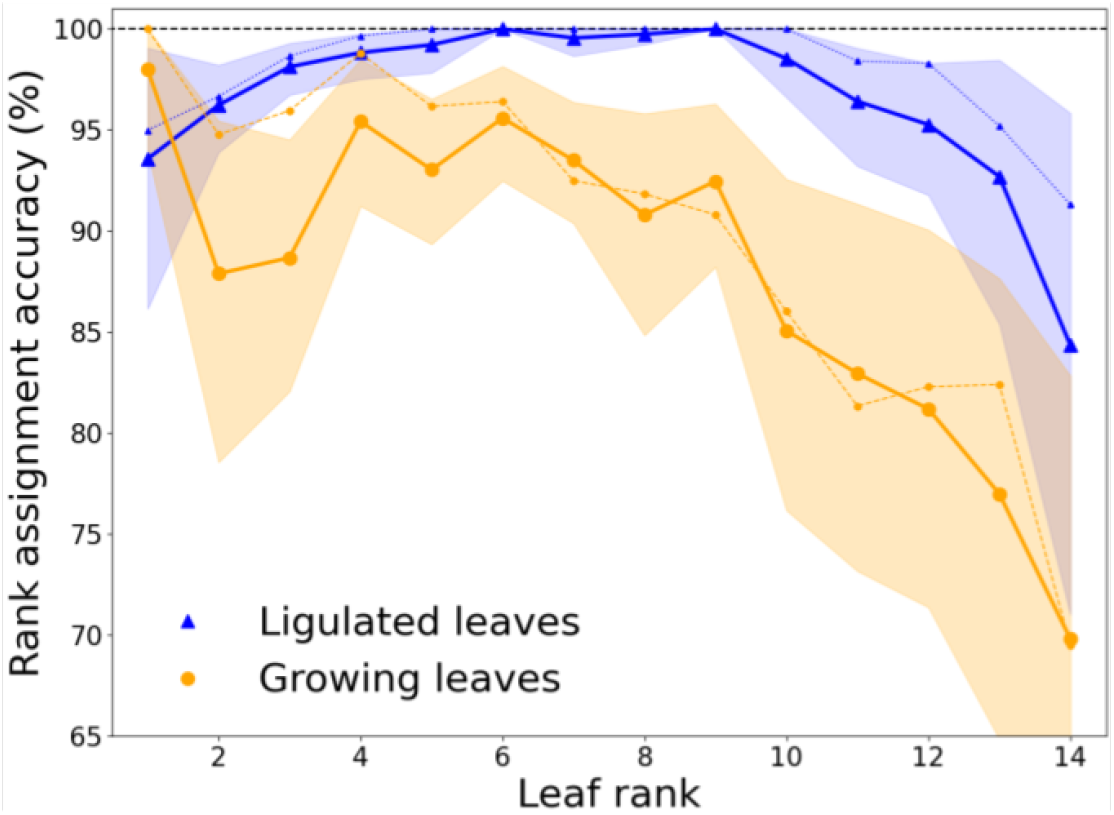
Accuracy of leaf rank assignment as a function of leaf rank. Leaf rank assignment accuracy is the percentage of non-artefact segmented leaves (n=10940) whose predicted rank matches ground-truth rank (observation). This metric is computed for each ground-truth rank value separately, for ligulated leaves (wide blue line) and growing leaves (wide orange line). Point value and error bars correspond respectively to the mean and 95% bootstrap confidence interval among 30 plants. Blue dotted line: ligulated leaves results without considering (i) leaf observations exceeding 20 day20°C after ligulation, and (ii) dates where maize ear was segmented as a leaf. Orange dotted line: growing leaves results after initialising their tracking with ground-truth ligulated leaf ranks.

Rank assignment accuracy was lower for growing leaves (85.3%, Table 1), but still high for the first bottom ranks (92,8% for ranks 1-10). This might be because a different algorithm is used compared to ligulated leaves, and because of the intrinsic difficulties associated with the detection of a developing organ: growing leaves cannot be topologically ordered *a priori*, and they may undergo rapid changes in shape and geometry over time. Also, growing leaves tracking used ligulated leaf ranks assignment as a starting point, which caused error propagation. Indeed, fewer errors were observed for lower leaves when manually initialising the growing leaf tracking with ground-truth ligulated leaf ranks (Fig. 6; orange dotted line). Overall, the errors caused by the growing leaf tracking are more frequent in the upper ranks, (Fig. 6; orange line) which might be due again to an increasing complexity of maize leaves structure over time. In particular, there are more leaves emerging at the same time in the maize whorl for late growth stages, and these leaves have a more similar morphology, making them difficult to distinguish (see Fig. 1).

### An automated quantification of plant development during the whole vegetative phase

This pipeline provides two ways to quantify plant development automatically: either (i) vertically, or (ii) through the number of emerged and ligulated leaves (leaf stage).

i. Stem height was predicted with high accuracy (RMSE = 2.02 cm, R^2^=0.999) for all growth stages. This provides a way to quantify plant vertical development regardless of how the leaves are deployed. This also provides a spatial delimitation of the mature and growing parts of the plant, which is more accurate than the morphological criteria used in [Artzet et al., 2019] (R^2^ = 0.68) and [Souza and Yang, 2021] (R^2^ = 0.92 for early growth stages), and remains robust in advanced stage once the maize ear emerges (Fig. 7A; no outliers for high observed values, i.e. advanced stages).
ii. The leaf stage predicted by the pipeline was correlated to the ground-truth observation (R^2^ = 0.87). Predictions were twice as accurate as when simply counting the leaves present on the reconstructed plant (Fig. 7B: RMSE = 1.29 vs RMSE = 2.62 for Phenomenal). Our method therefore avoids the bias that may occur in other leaf counting methods [Miao et al., 2021; Zhou et al., 2021; Souza and Yang, 2021] that do not take into account the disappearance of bottom leaves due to senescence. However, this trait was still consistently underestimated (bias = −1.09, Fig. 7B). The remaining error might be because the last leaves that have just emerged were often missing in the 3D reconstruction. This bias increases over time, since more and more leaves are growing simultaneously in the whorl [Ruget et al., 1996], as shown in Fig. 1. A linear regression can be applied to remove the bias from the prediction (red dashed line in Fig. 7B), which reduces the RMSE from 1.29 to 0.44.

**Fig. 7.**
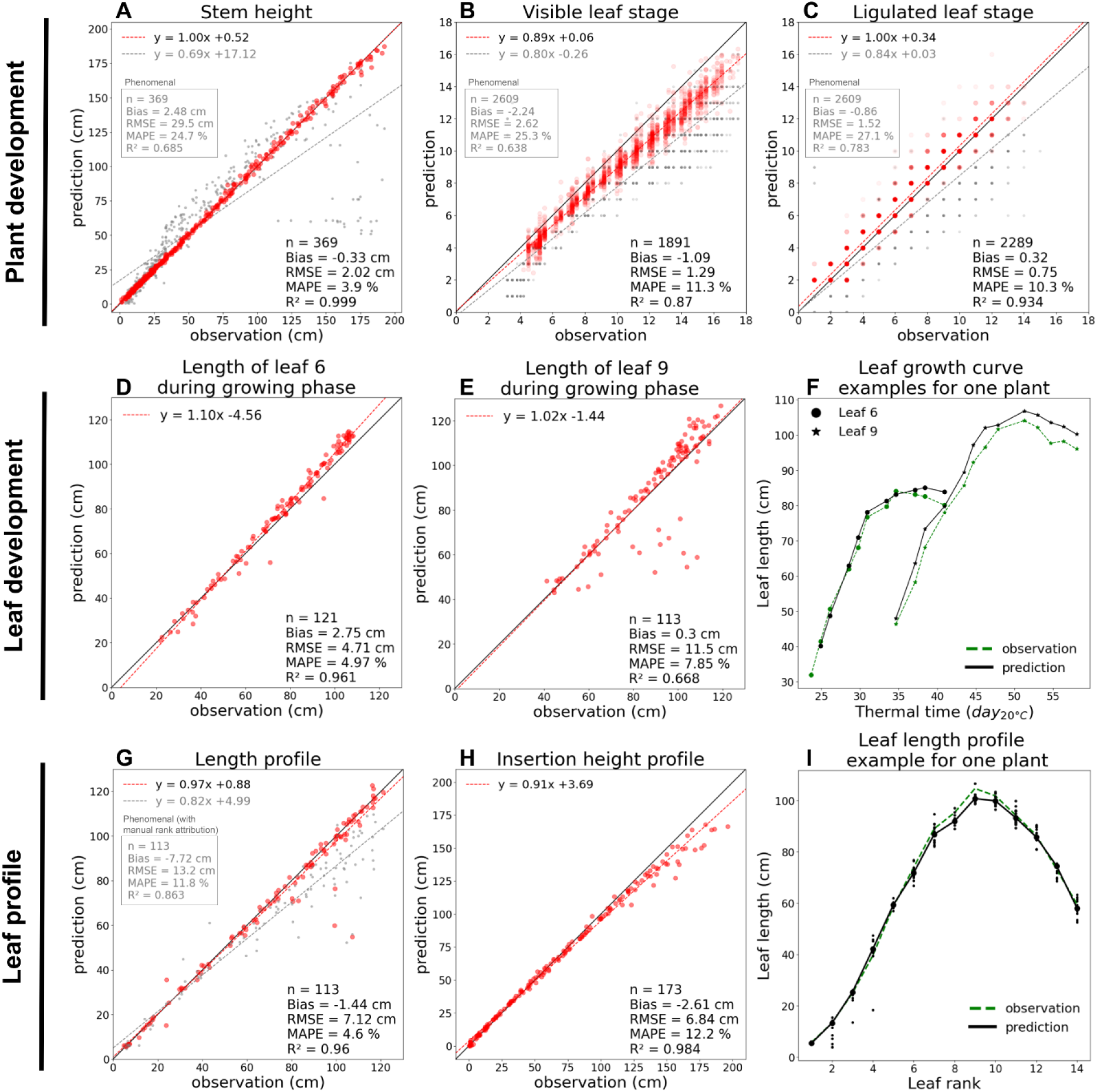
Evaluation of the phenotypic traits obtained with the pipeline. Pipeline predictions are compared with ground-truth observations for the following traits: **A)** Stem height, **B)** visible leaf stage, **C)** ligulated leaf stage, **D, E)** length of leaf 6 & 9 during growing phase, **G)** length profile, **H)** insertion height profile. For each trait, a linear regression is applied (x = observation, y = prediction). In **A, B, C, G**, the pipeline results are compared with Phenomenal [Artzet et al., 2019] outputs (points, regression equation and metrics are displayed in grey). Leaf growth (**G**) and Leaf length profile (**I**) outputs are illustrated for one representative plant. In **I**, larger points correspond to median predictions. n = number of points, RMSE = Root-Mean-Square Error, MAPE = Mean Absolute Percentage Error.

Leaf stage was also measured considering only the ligulated leaves, which resulted in a higher correlation (R^2^ = 0.93) and a lower bias (bias = 0.32, Fig. 7C). It is the first time that collar appearance rate, which is of crucial importance in maize models of development [Fournier and Andrieu, 1998; Lacube et al. 2020], can be measured with an automatic method at this degree of precision.

### An automated tracking of individual leaves development

The pipeline was used to automatically extract leaf growth dynamics for all leaf ranks. For evaluation, we focus on leaf length dynamics of leaves 6 and 9 before ligulation. For leaf 6, predictions were strongly correlated to ground-truth (R^2^ = 0.96, Fig. 7D). For leaf 9, the predicted length was close to ground-truth most of the time, but there were more outliers compared to leaf 6, resulting in a lower correlation (R^2^ = 0.67, Fig. 7E). Leaf dynamics seem to be less accurately captured for higher leaf ranks, which may be due to more frequent leaf rank assignment errors (Fig. 6, orange line), but also more generally because leaf lengths are less accurate for advanced stages due to frequent errors in the 3D organ segmentation (i.e. before tracking). Although the dynamics of leaf growth were only evaluated in terms of length here, the pipeline allows to capture the full evolution of leaf shape over time (Fig. 1), which can be described by other variables such as azimuth (Additional file 4A). Such leaf dynamics can also be extracted during the senescence phase when leaves collapse (Additional file 5).

### An automated reconstruction of plant architecture development

This pipeline was used to automatically extract leaf profiles, i.e. the estimation of various ligulated (i.e. mature but not senesced) leaf morphological features as a function of leaf rank. We considered leaf length and leaf insertion height profiles for evaluation, and both were highly correlated with ground-truth observations (length: R^2^ = 0.96, Fig. 7F. Insertion height: R^2^ = 0.98, Fig. 7G). As for leaf dynamics, the method can be extended to any other variable describing leaves, such as leaf width, internode width or leaf azimuth (Additional file 4B).

Thanks to tracking, median values can be derived from the successive measurements of morphological features for the same leaf over time. This significantly helps to minimise the errors that would have occurred using independent time points (e.g. ranks 2-4 in Fig. 7I). Outliers remaining after applying a median value were often observed for the last ranks (10 and more). This might be due to (i) more leaves overlapping in the upper part of the plant, therefore leading to less accurate 3D segmentations, (ii) the fact that the last leaves reach ligulation later and are therefore observed fewer times, which makes the calculation of a median less robust, (iii) a higher number of tracking errors for the last ranks.

## Discussion

### An adaptation of the sequence alignment framework to robust leaf tracking

Sequence alignment is proposed as an original solution to the leaf tracking problem, allowing to consider both (i) the topological information at a fixed date, by ordering the ligulated leaves in a sequence, and (ii) the redundancy of the geometric information over time, by describing each ligulated leaf as a vector of temporally invariant features. While sequence alignment has been applied outside the bioinformatics field [Abbott and Tsay, 2000; Prinzie and Van den Poel, 2006; Dieny et al., 2011], this is the first time, to our knowledge, it is used to track plant and organ growth. This framework allows us to consider the main difficulties of leaf tracking (segmentation artefacts, leaves appearance/disappearance) via the analogy of insertions and deletions of elements in a sequence. In plant phenotyping, tracking is often done step by step, by successive pairwise comparisons of reconstructed plant models, which could lead to the propagation of errors from the first alignments computed to the end of the time-series.

Sequence alignment methods allow the formulation of a global resolution algorithm for this optimization problem. In this study, we used an optimization method called progressive alignment [Batzoglou, 2005], which progressively integrates the models of successive time steps by comparing them to a “profile”, representing all the previous matched models. This method has shown great efficiency on our dataset.

Sequence alignment algorithms are known to be highly dependent on the choice of the parameters, especially the gap penalty [Notredame, 2002], thus further parameters fine tuning should be considered to optimise the method on more challenging datasets in the future. While this method was tested on maize plants, it could be adapted to any other species for which (i) the order of leaf emergence can be derived topologically along a single stem axis, and (ii) a subset of leaves with a stable geometry that can be identified at each time point (e.g. sunflowers or cotton). However, extending this method to branched plants, such as wheat or sorghum with tillers, remains an open problem.

### Temporal tracking enhances the robustness of 3D reconstruction

This work focused on the temporal processing of 3D segmented plant reconstructions. Each plant segmentation is performed at a given date independently of other dates, and can contain inaccurate leaf reconstructions, and segmentation artefacts. With our tracking method, potential error reconstructions can be compensated over time, by grouping several observations of a same leaf in the time-series. The sequence alignment framework also helps to overcome segmentation artefacts (e.g. missing leaves, ear misidentified as a leaf) by setting them apart. However, such segmentation errors were still often responsible for subsequent tracking errors in our dataset. This is particularly visible in the advanced stages of growth where the leaves emerge more frequently, are longer, and therefore intersect more, making the segmentation task more error-prone. It might therefore be better to spend time addressing these segmentation issues beforehand, rather than optimising the tracking parameters to address them later. For example, the maize ear could be detected beforehand in the segmented plant objects, using approaches similar to panicle [Brichet et al., 2017] and collar [Zhou et al., 2021] detection.

### A robust pipeline that can be used in high-throughput conditions

While other 3D+t phenotyping pipelines have already been proposed, they were mostly tested on a few number of plants and time points, for early growth stages (ca. 5-10 time points and 5-10 plants). Instead, the pipeline presented here was tested on a dataset of 355 plants of various genotypes grown under different environmental conditions. Each plant was observed through a time-series of ~43 time points covering all growth stages, making a total of 237,600 images analysed. Most traits were validated on a subset of only 30 plants due to time-consuming annotations, but leaf stages outputs were evaluated on the full dataset of 355 plants (see Fig. 7B-C). Other traits outputs are shown on Fig. 8 for the entire dataset of late harvesting plants, and overall exhibit coherent patterns for all ranks and growth stages. All these results suggest that this pipeline can be used in practice to phenotype large panels of maize plants.

**Fig. 8.**
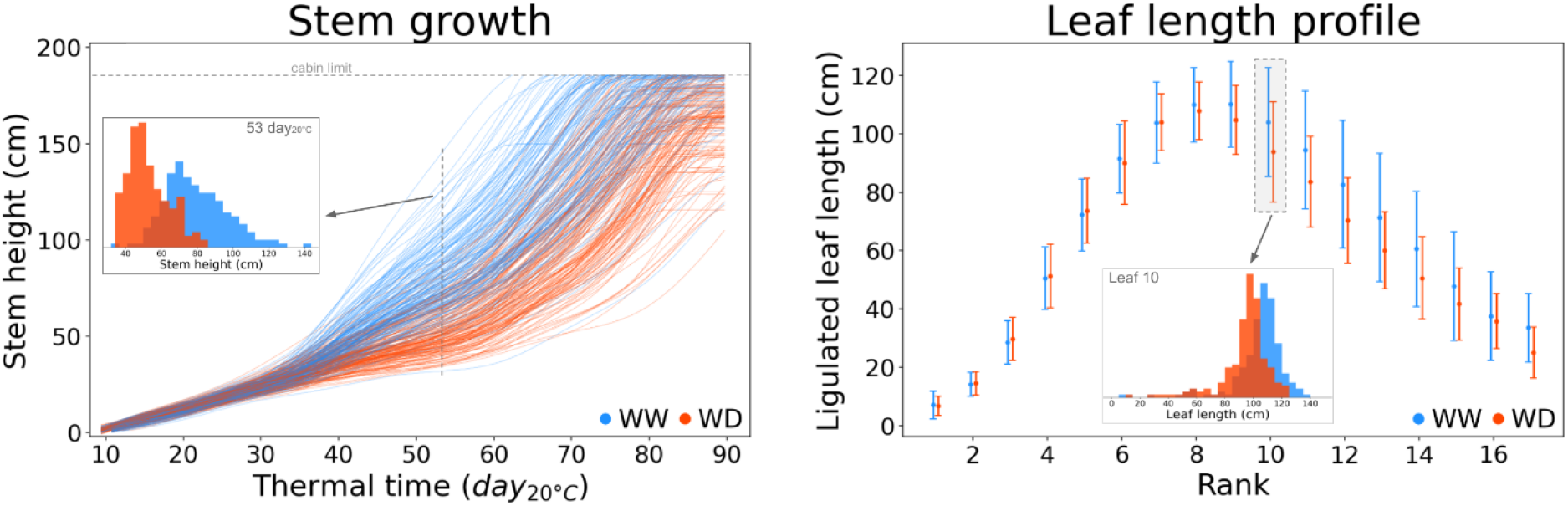
Automatic extraction of architecture and development traits at organ level in high-throughput conditions. Extraction of **A)** stem growth and **B)** leaf length profile using the pipeline, for 355 maize plants of 60 different genotypes, grown from plant emergence to flowering stage (237,600 images analysed), under well-watered (WW; 178 plants, blue) and water deficit (WD; 177 plants, red) conditions. **A**: one line per plant, each line is smoothed (see Step 5 of the pipeline). **B**: point = mean, error bar = standard deviation.

### Quantification of maize architecture and development for plant modelling

Plant development is usually quantified indirectly and incompletely in phenotyping platforms, using for example the height of the highest plant pixel on images [Golzarian et al., 2011] as a proxy of vertical development, or the total number of plant pixels [Cabrera-Bosquet et al., 2016]. On the contrary, our method directly measures detailed botanical features for all organs, together with their dynamics. For example, the stem height estimated with our pipeline fits the botanical definition of vertical plant growth. Other traits such as individual leaf elongation have already been measured semi-automatically in HTP platforms (e.g. transducer, [Sadok et al., 2007]), but such methods are limited to monitoring a small number of leaves of the plant, which can lead to serious limitation [Granier and Tardieu, 2009]. Instead, our method can quantify the growth dynamics of all leaves simultaneously, therefore capturing the full growth dynamics of the plant. Finally, traits such as leaf profiles are difficult to measure in a non-destructive way, and it has never been done on hundreds of maize plants to our knowledge.

Such automatic phenotypic measurements are valuable input data to parameterize models such as Functional-Structural Plant Models (FSPM) [Fournier and Andrieu, 1998; Cieslak et al., 2022], which fully capture the plant architecture and development up to the organ level. While these models are often calibrated via indirect proxies due to lack of additional available data, our 3D+t plant reconstruction method could give a more direct access to an accurate plant model calibration. Moreover, the phenotypic traits obtained with our pipeline show differences between different genotypes and watering treatments (Fig. 8), illustrating that the pipeline is sufficiently accurate to compare GxE interactions. Using our pipeline on large plant panels in HTP conditions could therefore provide the input data necessary to parameterize both genetic and environmental effects on plant architecture and development in FSPM models.

## Conclusion

The pipeline presented here allows reconstructing a time-consistent 3D architecture of a maize plant at organ level, from emergence to flowering. Leaf tracking is the main challenge in this kind of phenotypic analysis, and sequence alignment appears to be well suited for this task. Each detected leaf organ is labelled by its rank, which allows to fully describe and compare the architecture of the organs sharing the same position in different plants (e.g. leaves length). Moreover, the different observations of the same organ over time are grouped together, providing more accurate measurements by compensating eventual errors over time. The temporal consistency of the data makes it possible to describe the development of the plant, both at organ level (leaf growth dynamics) and plant scale (vertical growth, leaf stage). Since this pipeline is fully automatic, phenotypic traits can be measured on thousands of plants in HTP platforms, with sufficient accuracy to compare the development and architecture of various GxE interactions along the entire growth cycle.

## Supplementary information

**Additional file 1** Details on the training of the deep-learning model for maize collar detection. **Additional file 2** Video (.mp4) displaying leaf tracking for 3 maize plants.

**Additional file 3** Example of the anomaly detection step for one maize plant. **A)** two reconstructed plants removed during anomaly detection due to their abnormal stem shapes. **B)** Stem height smoothing over time, allowing to correct an outlier.

**Additional file 4** Example of azimuth traits extracted with the pipeline for one maize plant. **A)** Azimuth dynamics of individual leaves, up to 40 day20°C after their first detection. **B)** Leaf azimuth profile.

**Additional file 5** Visualisation of rank assignment following sequence alignment on a set of 3D ligulated leaf polylines. **A)** Visualisation of all ligulated leaves polylines in a time-series of 3D reconstructions of one plant. **B)** Assignment of leaf ranks on this set of polylines, using sequence alignment.

## Authors’ contributions

LCB supervised the experiment and acquired the data. BD designed the pipeline and analysed the data. CF, CP, LCB and RF provided advice on the conception of the pipeline. BD and RF wrote the manuscript and CF, CP and LCB reviewed and edited it. All the authors have approved the manuscript and have made all required statements and declarations.

## Acknowledgements

We are grateful to all members at the M3P platforms for providing technical support, conducting the experiments and collecting data.

## Competing interests

The authors declare that they have no competing interests.

## Availability of data and materials

The source code and examples are available on Github (https://github.com/openalea/phenotrack3d) under an Open Source licence (Cecill-C).

## Consent for publication

Not applicable.

## Ethics approval and consent to participate

Not applicable.

## Funding

This work was supported by the EU project H2020 731013 (EPPN^2020^).

## Notes

### Competing Interest Statement

The authors have declared no competing interest.

