## Additional file 1 for "PhenoTrack3D: an automatic high-throughput phenotyping pipeline to track maize organs over time"

A dataset of collar positions was created by annotating images of a random subset of 30 plants with various GxE interactions (excluding plants already used to validate the pipeline). For each plant, a set of 20 RGB images from various time points at various angles was selected. For each image, VGG Image Annotator [Dutta et al., 2019] was used to annotate the (x, y) position of each visible collar. Collars were only annotated when their position on the stem could be determined without ambiguity. A total of 3284 collars were annotated out of 600 images.

Then, for a given annotated image, a 416 x 416 subpart of the image (vignette) was selected, taking as its center a point randomly chosen along the stem. This vignette was attached to its corresponding labeled data, i.e. the position of the collars visible on the vignette. This process could be repeated automatically as many times as desired. Thus, 25 plants were used to generate a training dataset of 40,000 vignettes and labels. The 5 remaining plants were used to generate a validation dataset of 4,000 vignettes and labels. Half of the data was flipped horizontally to increase the diversity of the data.

The training dataset was used to train a Yolov4 object detection model. Training vignettes and labels correspond to the training input and output respectively. The Yolov4 model predicts bounding box positions (x, y, w, h) by default, where (x, y) is the position of the upper left box pixel, w the box width in pixels and h the box height in pixels. Since only the bounding box center position matters here, w and h were fixed at a constant value of 50 pixels in the training and validation datasets. The validation dataset was used for training monitoring. The yolov4-tiny configuration was used, and the training process was stopped after 8000 iterations.

[Dutta et al., 2019] Dutta A, Zisserman A. The VIA annotation software for images, audio and video. InProceedings of the 27th ACM international conference on multimedia 2019 Oct 15 (pp. 2276-2279).
