## Supplementary figures and images for "PhenoTrack3D: an automatic high-throughput phenotyping pipeline to track maize organs over time"

### Additional file 3

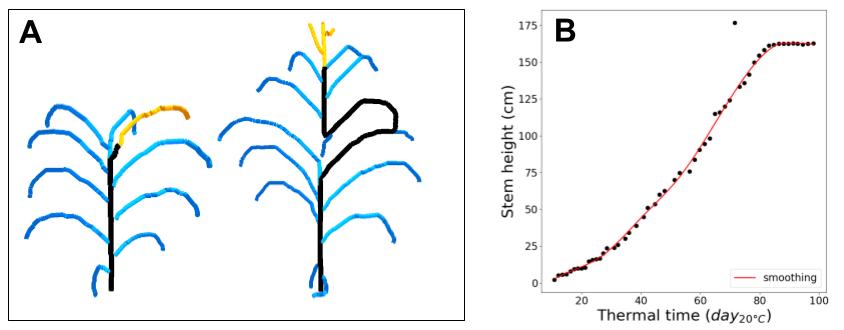

### Additional file 4

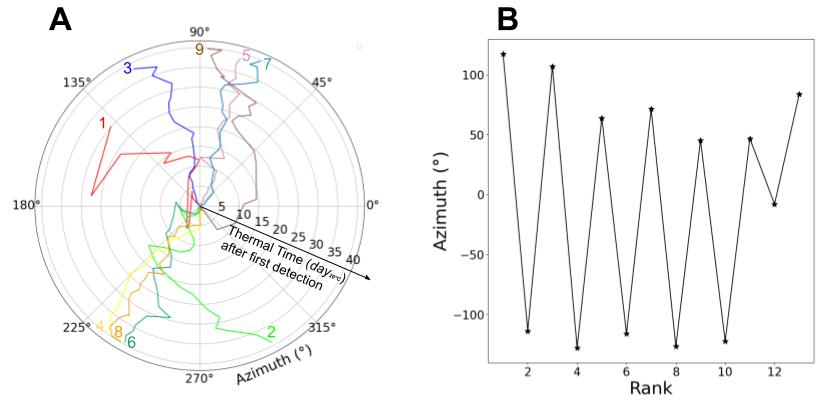

### Additional file 5

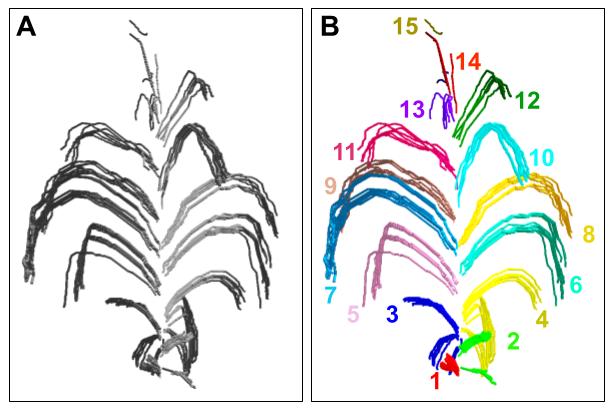
